# A Pre-Processing Pipeline to Quantify, Visualize and Reduce Technical Variation in Protein Microarray Studies

**DOI:** 10.1101/2021.09.29.461966

**Authors:** Sophie Bérubé, Tamaki Kobayashi, Amy Wesolowski, Douglas E. Norris, Ingo Ruczinski, William J. Moss, Thomas A. Louis

## Abstract

Technical variation, or variation from non-biological sources, is present in most laboratory assays. Correcting for this variation enables analysts to extract a biological signal that informs questions of interest. However, each assay has different sources and levels of technical variation and the choice of correction methods can impact downstream analyses. Compared to similar assays such as DNA microarrays, relatively few methods have been developed and evaluated for protein microarrays, a versatile tool for measuring levels of various proteins in serum samples. Here, we propose a pre-processing pipeline to correct for some common sources of technical variation in protein microarrays. The pipeline builds upon an existing normalization method by using controls to reduce technical variation. We evaluate our method using data from two protein microarray studies, and by simulation. We demonstrate that pre-processing choices impact the fluorescent-intensity based ranks of proteins, which in turn, impact downstream analysis.

**Impact Statement:** Protein microarrays are in wide use in cancer research, infectious disease diagnostics and biomarker identification. To inform research and practice in these and other fields, technical variation must be corrected using normalization and pre-processing. Current protein microarray studies use a variety of normalization methods, many of which were developed for DNA microarrays, and therefore are based on assumptions and data that are not ideal for protein microarrays. To address this issue, we develop, evaluate, and implement a pre-processing pipeline that corrects for technical variation in protein microarrays. We show that pre-processing and normalization directly impact the validity of downstream analysis, and protein-specific approaches are essential.

---

Protein microarrays are used to measure quantities of antibodies and other proteins in sera. They are versatile and are available for use in various fields such as cancer research, infectious disease diagnostics and biomarker identification, ^[5,10,16]^. Production of protein microarrays can occur in both commercial and academic laboratories and these varying laboratory conditions induce a wide range of technical effects. Effective characterization and adjustment for technical effects is necessary to ensure that inferences from protein arrays, to the degree possible, are based on biological variation. Normalization methods for similar assays, such as DNA arrays do not address the variable binding affinities and high sample-to-sample variability unique to proteins ^[3]^. Currently, investigators use a wide variety of normalization methods and pipelines^[14,15]^, but only a few protein microarray-specific methods are available ^[4,11,12]^. The availability of protein-specific methods, enables analysts to directly answer biological questions of interest using protein microarray data, many of which are addressed by ranking proteins based on their fluorescent intensity, then using these ranks to select candidate proteins that may be associated with observable phenotypes^[7,13]^. Proteins with the lowest fluorescent intensities are often excluded from analysis, therefore normalization methods can impact downstream analysis by changing ranks of proteins and altering which proteins are included in a final analysis ^[2,8]^.

We report on a pre-processing pipeline that leverages information from assay controls to improve upon existing normalization methods. We envision it as preparing data for input to subsequent modeling, for example, a Bayesian hierarchical model that extracts true signal from measured intensities. We measure sources of technical variation in two previously published protein microarray studies^[6,9]^using Bland-Altman plots^[11]^and associated regression analysis to evaluate the degree to which each pipeline step corrects technical variation. We identify sources of technical variation and data features that should be taken into account and others that normalization may not be able to correct, but could be accommodated in subsequent analyses. Using simulation, we show that the pipeline reduces technical variation while preserving true underlying signals in arrays that have technical variation and in those that do not. Finally, we demonstrate that the global ranks of protein levels are sensitive to which normalization methods are employed.

We implement the pre-processing pipeline in R version 4.1.0 and use functions in the limma package. The full code to reproduce all analysis in this manuscript can be found on GitHub (https://github.com/sberube3/pre_processing_proteomics). The pre-processing pipeline steps are: (1)log transformation, (2) removal of array and sub-array level effects with a robust linear model (RLM), and (3) standardization using control probe values. The full details are in Section S1. A regression analysis of Bland-Altman plot inputs evaluates the level of measurement agreement within and between arrays at each stage of the pipeline (see Section S2 for details). Specifically, we evaluate the level of measurement agreement between duplicated protein spots on a single array to assess within-array technical variation, and evaluate the level of measurement agreement between spots that measure the same protein on different control serum arrays to assess between-array technical variation. We then compare the ranks of proteins at each stage of the normalization pipeline. Section S4 gives details on ranking, and Section S5 describes the samples and protein microarrays used in each study. Finally, Section S6 describes the simulation setup.

We evaluate our proposed pipeline on data from two protein microarray studies, one on the identification biomarkers in early stages of lung cancer which uses HuProt^*™*^arrays and one on the antibodies associated with malaria exposure which uses homemade antibody arrays (full details on the study set up and arrays used in Section S5). Specifically, using three HuProt^*™*^arrays^[9]^and three malaria arrays^[6]^, we observe that the slopes and intercepts of OLS lines in Figure 1 are non-zero and conclude that every array has residual technical variation after RLM normalization. However, the modified Bland-Altman plot has a different slope and intercept for each array, suggesting that they may have a different level of residual technical variation ^[12]^.

**Figure 1:**
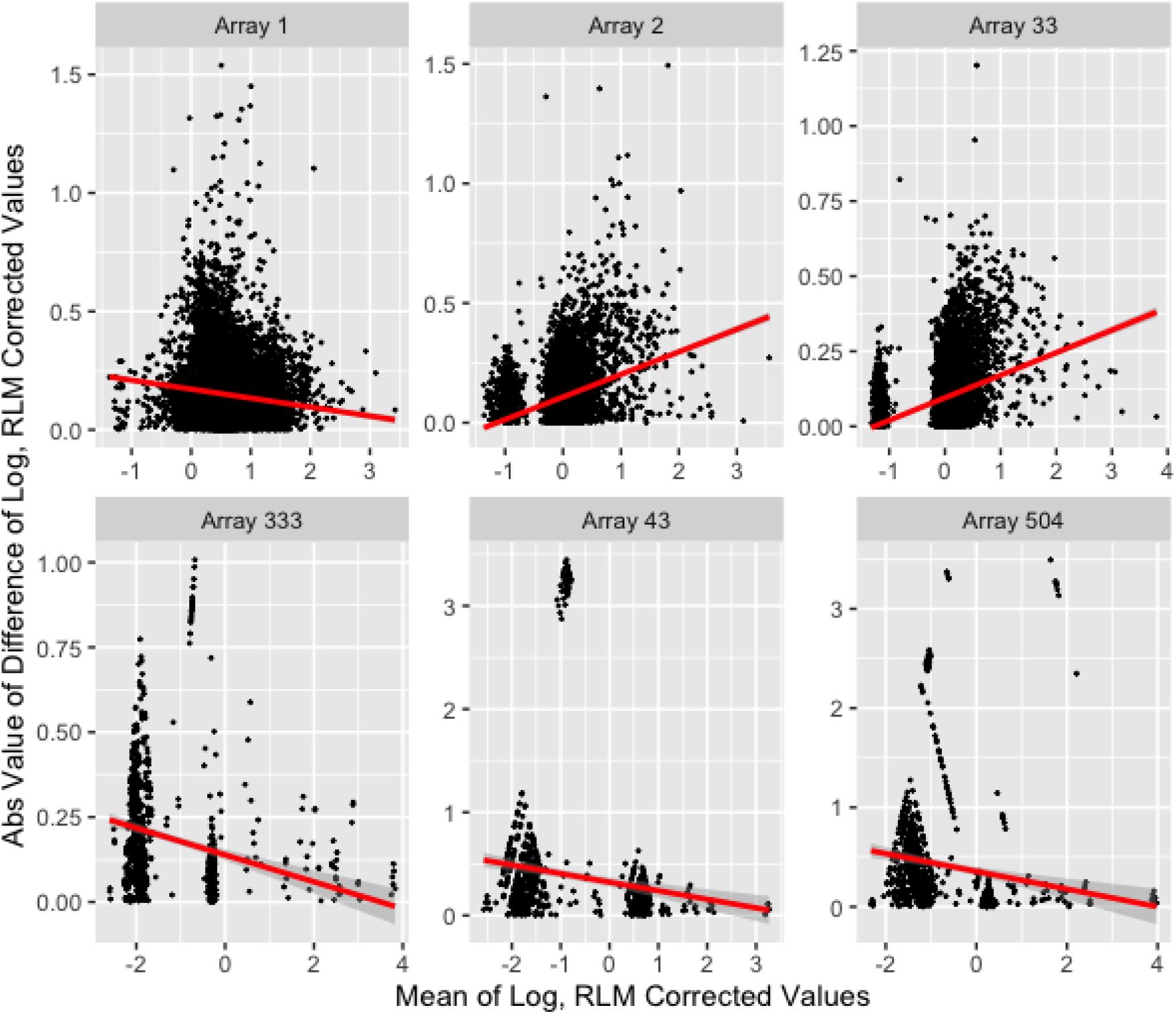
Bland-Altman Plots of HuProt^*™*^ arrays^[9]^ (Arrays 1, 2, 33) and malaria arrays^[6]^ (Arrays 333, 43, 504). We show *M* vs |*D*| with 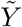s of duplicated probes from a single array. The ordinary least square line is in red. Outlying pairs of probes have been removed.

Using pairwise comparisons across control sera from the HuProt^*™*^and malaria arrays, we observe a lack of measurement agreement after RLM normalization in Figure S3 and Table S3. Specifically, slopes and intercepts of OLS regression for all arrays under RLM normalization are non zero and concordance correlation coefficients 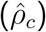 range from 0.117 to 0.618 in the malaria arrays and 0.240 to 0.841 in the HuProt^*™*^ arrays. This finding provides evidence that an additional pre-processing step is necessary. After standardization, which is aimed at homogenizing both the mean and variance of RLM corrected control probes across arrays, we observe increased measurement agreement between control serum arrays through the higher concordance correlation coefficients 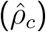 in Table S3 of the standardized measurements. In Section S7 we evaluate other methods of standardizations and further break down the goals of homogenizing the mean and variance of intensities across control sample arrays. We conclude that the traditional method of standardization with mean and variance outperforms other methods of standardization.

To further compare RLM normalization alone to RLM normalization with subsequent standardization, we perform a simulation to evaluate performance of the pipeline under four different scenarios. In the case where there is no technical variation between arrays (Scenario 1), there is little difference between standardization and RLM correction, when there is technical variation across arrays (Scenario 2), standardization substantially increased measurement agreement between arrays, and preserves true differences between measurements (mean difference of the 12 spike-in controls across arrays is close to 0). Furthermore, when the log transformation is inappropriate (Scenarios 3 and 4), the standardized values show strong measurement agreement, evidenced by the near zero mean squared error for differences of means and variances (Table 1), and preserves true differences between arrays, something RLM correction alone does not accomplish, most markedly when there is both model misspecification and technical variation between arrays (Scenario 4).

**Table 1:** Mean squared error (MSE) of difference in means and variances across pairs of arrays, and mean differences of spike-in controls across arrays. Sample mean of 1000 simulations and, in subscript, the sample standard deviation.

| Array Type | Scenario | Method | MSE Mean | MSE Var | Mean Difference |
| --- | --- | --- | --- | --- | --- |
| Malaria | 1 | RLM | 0.013<sub>0.009</sub> | 0.017<sub>0.013</sub> | -0.001<sub>0.098</sub> |
|  |  | RLM+ Std. | 0.003<sub>0.002</sub> | 0.006<sub>0.004</sub> | 0.001<sub>0.033</sub> |
|  | 2 | RLM | 157.760<sub>5.865</sub> | 1.145<sub>0.162</sub> | 4.357<sub>0.128</sub> |
|  |  | RLM+Std. | 0.015<sub>0.006</sub> | 0.296<sub>0.047</sub> | 0.357<sub>0.035</sub> |
|  | 3 | RLM | 1.840<sub>1.353</sub> | 0.282<sub>0.200</sub> | -0.010<sub>1.367</sub> |
|  |  | RLM+ Std. | 0.002<sub>0.001</sub> | 0.005<sub>0.004</sub> | -0.001<sub>0.037</sub> |
|  | 4 | RLM | 2739.451<sub>122.545</sub> | 0.289<sub>0.212</sub> | 70.966<sub>3.276</sub> |
|  |  | RLM+Std. | 0.018<sub>0.006</sub> | 0.022<sub>0.012</sub> | 0.291<sub>0.037</sub> |
| HuProt<sup>TM</sup> | 1 | RLM | 0.000<sub>0.000</sub> | 0.001<sub>0.001</sub> | -0.000<sub>0.012</sub> |
|  |  | RLM+Std. | 0.001<sub>0.000</sub> | 0.022<sub>0.015</sub> | -0.000<sub>0.014</sub> |
|  | 2 | RLM | 2.580<sub>0.068</sub> | 0.588<sub>0.028</sub> | 1.072<sub>0.017</sub> |
|  |  | RLM+Std. | 0.029<sub>0.004</sub> | 3.358<sub>0.317</sub> | 0.772<sub>0.017</sub> |
|  | 3 | RLM | 0.002<sub>0.001</sub> | 0.016<sub>0.011</sub> | 0.003<sub>0.039</sub> |
|  |  | RLM+Std. | 0.000<sub>0.000</sub> | 0.012<sub>0.008</sub> | 0.000<sub>0.018</sub> |
|  | 4 | RLM | 17.377<sub>0.362</sub> | 0.026<sub>0.017</sub> | 3.406<sub>0.064</sub> |
|  |  | RLM+Std. | 0.175<sub>0.011</sub> | 1.748<sub>0.256</sub> | 0.822<sub>0.037</sub> |

Using the simulation setup (Section S6) we evaluate three other common methods of normalization: quantile, variance stabilizing and Cyclic Loess^[3]^. We conclude that while certain metrics in some scenarios suggest that these methods outperform RLM normalization followed by standardization, in most instances these methods produce highly similar results (Table S6). Importantly however, these three methods, outperform RLM normalization alone in all scenarios, further hilighting the added value of standardization. Additionally, the distribution of fluorescent intensities from arrays with serum from malaria endemic countries and from individuals with cancer (non-control sera arrays), in both datasets (non-simulated) after normalization reveal that much of the between array variability present in the raw data, some of which is biological, is erased by quantile normalization, variance stabilizing normalization, and Cyclic Loess normalization. By contrast, RLM normalization followed by standardization preserves much of this variability (Figure S9) suggesting that RLM normalization followed by standardization is the best option for removing technical variation while preserving true underlying biological signal.

Given that ranking is a common analysis method for protein microarray data, in order to assess the impact of pre-processing steps on downstream analysis, we compare ranks of proteins by fluorescent intensity under different normalization procedures. For both the malaria and HuProt^*™*^ arrays, the ordering of the top 10 proteins shows little dependence on normalization method. In contrast, when considering the top 25 − 35 proteins, there are several changes in ordering at each stage of normalization, changes that could impact which proteins are targeted for analysis, and that underscore different proteins of interest than the ones previously identified by Kobayashi et al.^[6]^(Figure 2). The 170 proteins identified by Pan et al.^[9]^ were not identified by ranking, however, none of the proteins identified in this publication overlap with the top 30 at any normalization step.

**Figure 2:**
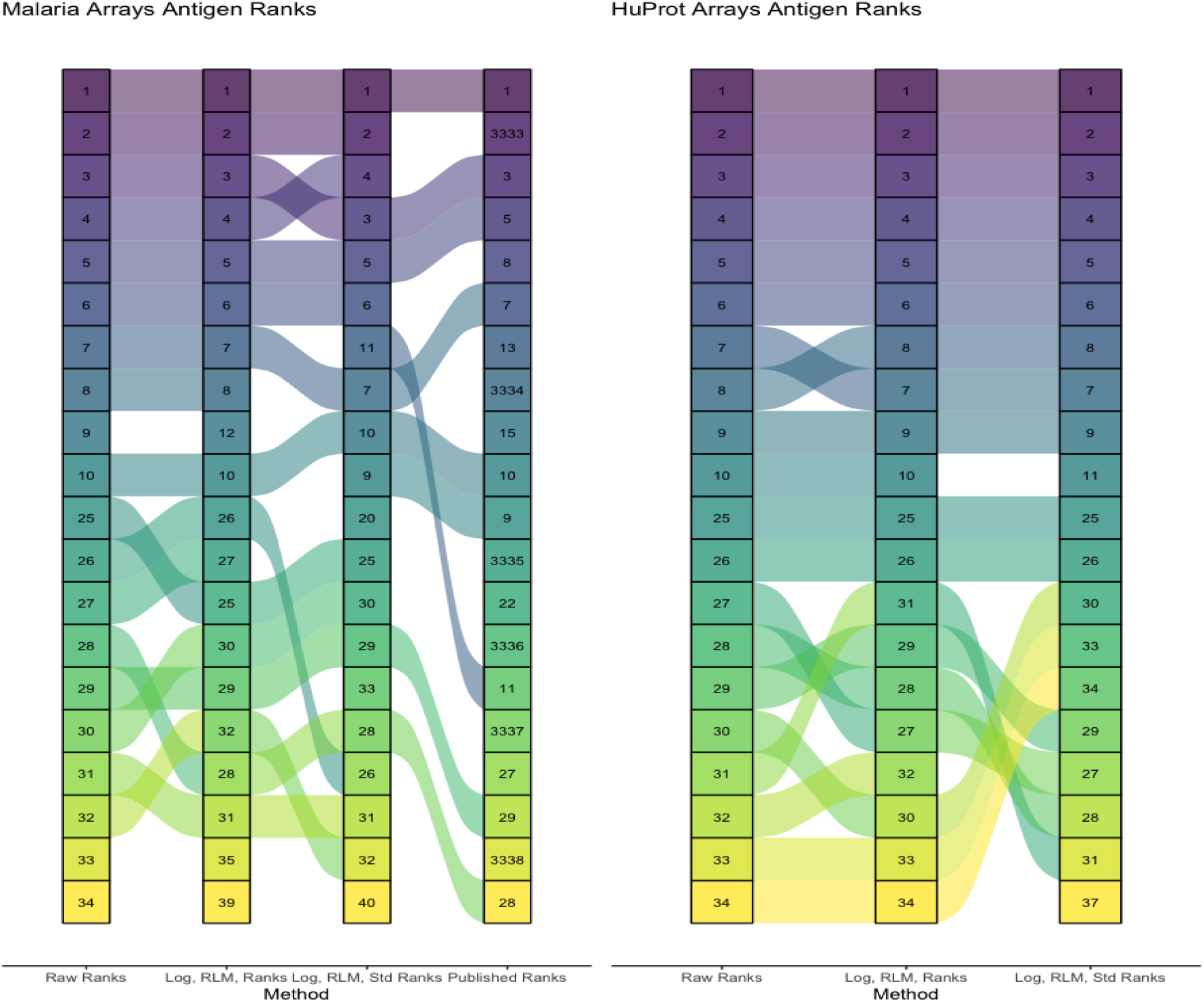
Ranks of top 10 and top 25 − 35 antigens on malaria and HuProt^*T M*^ (cancer) arrays are compared under three different conditions: raw intensities (*Y*′), log-transformed and RLM corrected intensities 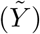 and log-transformed, RLM corrected and standardized intensities (*Y*)all as described in Section S1. Additionally the published ranks from Kobayashi et al.^[6]^ are added to the malaria array ranks. Proteins that are not published in the top 30, are represented with the placeholders 3333 *-* 3338.

Performing effective normalization on protein microarray data sets the stage for analyses that can directly answer questions of biological interest. Using Bland-Altman plots, associated regression analyses, and concordance correlations we have evaluated a pipeline for normalizing measurements from protein microarrays. We show that within-array and between-array technical variability are present in both protein microarray studies after RLM normalization^[12]^. Standardization conducted using information from control probes reduces some between-array technical variation in both the malaria and HuProt^*™*^, this finding is further supported by results of a simulation study.

Even after standardization, technical variation within and across arrays remains to varying degrees. Flexible model-based normalization approaches are the best candidates to address these differences. These normalization approaches are likely to have the greatest impact on data from arrays that are manufactured on a small scale by academic labs. However, given the differences in the number of replicated probes and the target proteins between the HuProt^*™*^ and malaria arrays, we cannot directly compare the impact of our pipeline on the results of the two studies. Furthermore, even flexible normalization procedures that take into account different levels and characteristics of technical variation within-and between arrays are unlikely to fully account for some extreme outliers both at the array and probe levels. Therefore, identifying these outliers through Bland-Altman analysis and setting them aside is likely the wisest course of action. We evaluated our normalization procedures for two published studies and one simulated dataset, representing a fairly narrow sample of the currently available protein microarrays. We evaluate the ability of our pipeline to correct technical variation while preserving true biological signal with *simulated* spike in control probes, however, a laboratory experiment with spike in probes is a more direct way to evaluate this.

The goal of normalization of a laboratory assay is to remove technical variation in order to focus inferences on biological processes. Understanding how technical variation manifests in an assay is therefore a key step in designing a normalization procedure that corrects for this variation. Given the versatility and the variety of manufacturing processes for protein microarrays, a systematic and quantitative investigation of technical variation across studies is relevant for developing broadly usable analysis pipelines. This is an important investigation especially in light of the sensitivity of protein ranks to the normalization steps employed. Adding standardization to existing methods corrects some technical variation, but further normalization models that incorporate features of the data revealed by our Bland-Almtan analysis, such as edge effects, are necessary and will likely have an important impact on biological inference.

## 2 Acknowledgments

The authors are grateful to Philip Felgner and D.Huw Davies for sharing data relating to their *Plasmodium falciparum* and *P. vivax* antibody arrays. The authors also acknowledge Jianbo Pan and Heng Zhu for sharing the protein microarray data from their lung cancer study.

