## Supplement for "A Pre-Processing Pipeline to Quantify, Visualize and Reduce Technical Variation in Protein Microarray Studies"

July 13, 2021

#### 1 The Pre-Processing Pipeline

##### 1.1 Step 1: Log-Transformed Readings

Let $R_{ai}$ be the measured fluorescent intensity (the reading) from source "a" and $R_{bi}$ the reading from source "b" with "$i$" indicating pairing. The inference related to Bland-Altman plots relies on the assumption that the vertical axis $(R_{ai} - R_{bi})$, the difference of the paired readings, is normally distributed,[1] or at least symmetric and unimodal with no edge effects. But, the R-values are non-negative and so we use natural log-transform values: $Y'_{ai} = \log(R_{ai})$.

##### 1.2 Step 2: Robust Linear Model for Normalization

In order to account for probe-level, sub-array level and array level fixed effects, Sboner et al.[9] suggest a linear model with robust approach (using an M-estimate) to estimate these effects and use model residuals as inputs for downstream analysis. We follow that approach, expand the subscripts, and apply the linear model:

$$Y'_{ijkr} = \alpha_i + \beta_j + \tau_k + \epsilon_{ijkr} \tag{1}$$

to control probes only (all positive and negative control probes in Tables 1 and 2), where $Y'_{ijkr}$ is the log-transformed measured fluorescent intensity (the reading) on array $i$, in sub-array $j$ for replicate $r$ (generally, at most 2) of probe $k$. Therefore, $\alpha_i$ is the effect for array $i$, $\beta_j$ the effect for sub-array $j$, $\tau_k$ the probe effect for protein $k$, and $\epsilon_{ikjr}$ is the spot specific unexplained variation for the control probes, assumed to be approximately Gaussian distributed with mean $0$ and variance that doesn't depend on subscripts.

Estimates of $\alpha, \beta$ and $\tau$ are obtained by fitting the model to the set of control probes $(Y'_{ictrl})$ across all arrays and sub-arrays in a study. That is, all array, sub-array and probe level effects are jointly estimated for a study. We then compute residuals for all data (control and active probes), producing,

$$\tilde{Y}_{ijkr} = Y'_{ijkr} - \hat{\alpha}_i - \hat{\beta}_j \tag{2}$$

and use the values $\tilde{Y}_{ijkr}$, with array and sub-array effects removed, in subsequent analyses. While we include $\tau_k$ in the model (Equation 1) to ensure that $\hat{\alpha}_i$ and $\hat{\beta}_i$ are adjusted for probe level effects, given that the model is only fit with control probes, it would not be appropriate to subtract these control probe-level effects from all test probes, therefore we only subtract $\hat{\alpha}_i$ and $\hat{\beta}_i$ to obtain $\tilde{Y}_{ijkr}$ (Equation 2).

##### 1.3 Step 3: Standardization

Sets of control probes are often comprised of blank spots and proteins that are known to be ubiquitously present at high concentrations in the samples being studied. Therefore, regardless of the sample's phenotype, we anticipate strong measurement agreement of control probes across arrays if technical variation has essentially been removed. After removal of array level-effects described in Section 1.2, the sample mean over the set of RLM corrected control probes on array $i$, $\tilde{Y}_{i,ctrl}$, should be approximately zero (because the model is estimated robustly, it won't be exactly zero). However, the sample variances of the $\tilde{Y}_{i,ctrl}$ may still be different. Therefore, we apply and evaluate a correction to ensure that the sample variance of all corrected control probes $\tilde{Y}_{ctrl}$, across all arrays is nearly constant. Specifically, we let,

$$Y_{ijkr} = \frac{\tilde{Y}_{ijkr} - \hat{\mu}_i}{\hat{\sigma}_i}, \tag{3}$$

where $\hat{\mu}_i$ is the sample mean over the set of RLM corrected control probes (all positive and negative control probes in Tables 1 and 2) from array $i$, $\tilde{Y}_{i,ctrl}$, and $\hat{\sigma}_i$ is the sample standard deviation over this same set. We evaluate two other methods of standardization: one with the sample median and the "pseudo-variance" (inter-quartile range (IQR) divided by $1.35$) ; the other using the 10% (5% on each end) Winsorized mean and sample standard deviation. Note that while $\hat{\mu}_i$ and $\hat{\sigma}_i$ are computed using only control probe intensities, the correction in equation 3 is applied to all probes.

We also evaluate a Box-Cox transform as an alternative to the log transform, however, in most cases the best transform is close to the logarithm ( Figure 2).

Box-Cox transform of raw measurements, $R$, $\frac{R^{\lambda}-1}{\lambda}$ for $\lambda \neq 0$, and $\log(R)$ for $\lambda = 0$. We estimate $\lambda$ separately for each array in each study using a maximum likelihood approach.

#### 2 Bland-Altman Analysis

In order to characterize measurement agreement across protein microarrays, we use observed signals on the array described in Section 5. Specifically, $R_{ai}$ is the ratio of observed foreground to observed background of a probe $i$ from source $a$ and $R_{bi}$ is the ratio of observed foreground to observed background of this same probe from source $b$.

We consider the following notation for each ordered pair on a Bland-Altman plot where values can be raw intensities, log-intensities or corrected log-intensities (see Section 1):

$$\left(M_i = \frac{R_{ai} + R_{bi}}{2}, D_i = R_{ai} - R_{bi}\right), \tag{4}$$

In order to understand the utility of the Bland-Altman plot, we compute the relationship between $M$ and $D$ via the Ordinary Least Squares (OLS) fit obtained by regressing $D$ on $M$. With $R_a$

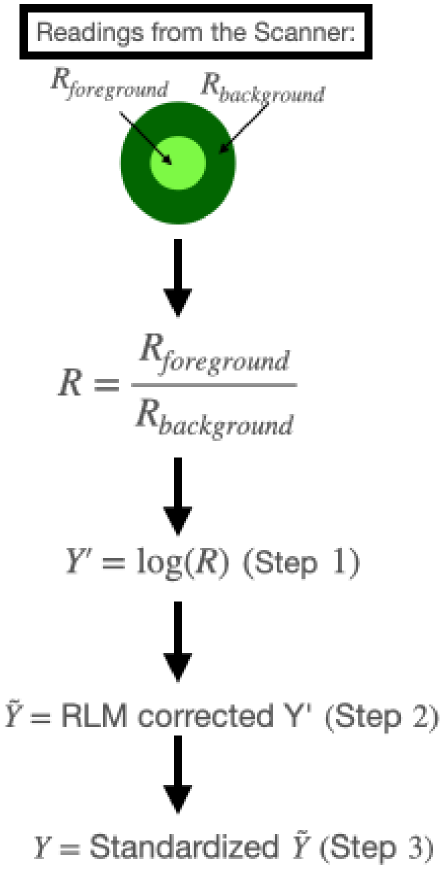


**Fig S 1:** Schematic of pre-processing pipeline, starting with values obtained directly from the protein microarray scanner.

and $R_b$ the measurements for sources $a$ and $b$, $s_a^2$ and $s_b^2$ the sample variances for $R_a$ and $R_b$, $\text{Cov}(R_a, R_b)$ the sample covariance of $R_a$ and $R_b$ and $\bar{M}$ and $\bar{D}$ the means of $M$ and $D$ from equation 4, the slope and intercept are:

$$\hat{\beta}_1 = \frac{2(s_a^2 - s_b^2)}{s_a^2 + 2\text{Cov}(R_a, R_b) + s_b^2}, \quad \hat{\beta}_0 = \bar{D} - \hat{\beta}_1 \bar{M} \tag{5}$$

Thus, the $\hat{\beta}_1$ measures the difference in variability across the two sources of measurement relative to the variability of the sum of measurements from the two sources. In the case of perfect agreement across both sources of measurement, the difference in variability is zero and thus the slope of the OLS line is also zero. More realistically, measurement agreement across sources in an assay will not be absolute, but slopes close to zero can still be observed indicating strong measurement agreement (see Bland and Altman p. 309 for an example of this[2]).

Since $\hat{\beta}_0$ is a function of both slope and the mean difference of $R_a$ and $R_b$, this parameter gives the analyst information about both differences in variability, and average difference in magnitude of measurements from sources $a$ and $b$. With $\hat{\beta}_1 = 0$, $\hat{\beta}_0$ is exactly the average difference in measurement magnitude between source $a$ and $b$.

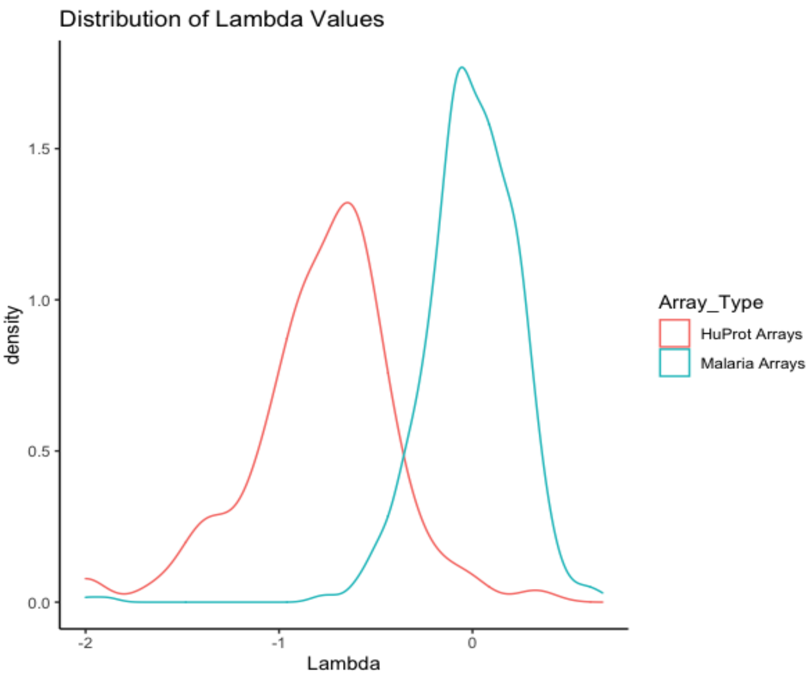


**Fig S 2:** The distribution of lambda values for the Box-Cox transform of the readings ($R$) estimated separately for each array in each study using a maximum likelihood approach.

The OLS line provides information about the linear relationship between $M$ and $D$, however, the shape of points around this line also provides insight about measurement agreement. Specifically, the association between the magnitude of measurements and their variability can be inferred by observing the shape of points on the Bland-Altman plot. In cases where the magnitude and the standard deviation have a strong positive correlation, points should fan away from the OLS line as $M$ increases. Additionally, points in a diamond shape around the OLS line suggest an upper and lower bound on the difference between measurements. We described how transformations of $R, M$ and $D$ can impact the shape of points around the OLS line in Section 1.

Bland-Altman plots can also diagnose outlying sources of measurements through multiple pairwise comparisons across sources of measurement either at a cursory glance or by analyzing residuals of a simple linear regression of $D$ on $M$.

#### 3 Sources of Technical Variation

We consider two main sources of technical variation in evaluating the pipeline: within and between array technical variability. In both instances, duplicated probes between (control probes) and within (replicated probes) arrays can be used to assess and correct technical variation.

Within array technical variability arises when laboratory conditions, such as solvent concentration, differ within sub-regions of a single array (sub-array effects), and is measured by observing differences across replicated probes on a single array. Therefore Bland-Atlman plots aimed at measuring within array technical variability use as inputs replicated probes within an array:

$$\left(M_i = \frac{R_{1i} + R_{2i}}{2}, D_i = R_{1i} - R_{2i}\right), \tag{6}$$

where $i$ refers to the array index, and $1$ is the first replicate of a particular probe, and $2$ is the second replicate of that same probe. In the HuProt$^{TM}$ arrays, we use all proteins on the array in the Bland-Altman analysis, since all proteins are present in duplicate on each array. In the malaria arrays we only use control proteins in the Bland-Altman analysis since only these are present more than once on each array. In addition to within-array technical variability, elements of the laboratory procedure can induce between array technical variability. Analyzing differences across arrays that are spotted with serum intended to be non-reactive to the probes on arrays, or control serum, allows us to characterize between-array technical variation. It is standard to include arrays with control serum in a study, such as cancer-free individuals in the cancer study and individuals residing in non-malarious regions in the malaria study. Given that we expect measurements from control serum samples to have high degrees of measurement agreement in the absence of technical variability, characterizing measurement agreement across the same probe on different control serum arrays, reveals between array technical variation. Bland-Atlman plots aimed at measuring between array technical variability used as inputs

$$\left(M_i = \frac{R_{li} + R_{1i'}}{2}, D_i = R_{li} - R_{1i'}\right), \tag{7}$$

where $i$ and $i'$ are the indices of two different arrays, and $l$ is the index of a particular probe at a particular location on the array, typically the layout of probes do not change from array to array within a study. We use all proteins (control and active) in Bland-Altman plots for both the HuProt$^{TM}$ and malaria arrays.

### 4 Protein Ranks

Central to developing broadly applicable normalization techniques is an evaluation of the impact of pre-processing procedures on downstream analyses. A common component of protein microarray analysis is identifying a pared down list of target proteins, often the $30$ to $50$ proteins with the highest fluorescent intensities, to directly compare across observable phenotypes, such as disease status. Identifying target proteins for analysis based on ranking fluorescent intensities is often done after normalization, but ranking procedures are sensitive to which normalization techniques the analyst applied. For both example datasets, we consider the median fluorescent intensity for each probe across all arrays, these median probe-level intensities are then ranked as point estimates.

### 5 Protein Microarray Data

The HuProt$^{TM}$ (v. 3.0) arrays are functional protein microarrays manufactured by CDI labs that contain 43776 test probes corresponding to approximately 16000 different proteins in the human proteome arranged into 24 blocks (or sub-arrays) on each slide. The arrays also contain 4608 control probes corresponding to multiple replicates of negative controls such as empty spots as

**Table S 1:** Probe composition on the HuProt$^{TM}$ array.

| Protein Category | Protein | Number of Spots |
|---|---|---|
| Negative Control | GST 10 ng/$\mu$l | 48 |
| | GST 50 ng/ $\mu$l | 48 |
| | GST 100 ng/$\mu$l | 48 |
| | GST 200 ng/$\mu$l | 48 |
| | Mouse-anti-biotin | 48 |
| | Rabbit-anti-biotin | 48 |
| | BSA | 48 |
| | Buffer | 816 |
| | Empty | 3072 |
| Positive Control | Histone 1 | 48 |
| | Histone 2 (A+B) | 48 |
| | Histone 3 | 48 |
| | Histone 4 | 48 |
| | Alexa Fluor labeled IgG | 48 |
| | Rhodamine+ Alexa Fluor labeled IgG | 48 |
| | Biotin-BSA | 48 |
| | Mouse IgM | 48 |
| Active Proteins | Active Proteins | 48384 |

well as positive controls comprising proteins to which antibodies are commonly found at high levels in human serum (Table 1). This study analyzed the antibody profiles of $100$ individuals. Of these individuals, $20$ were healthy (control serum) and the remaining $80$ had a form of lung cancer.[8] Further information on the protocols used to construct, probe and analyze these arrays can be found in Hu et al.[6], Hu et al.[5], and Yang et al.[10].

The malaria arrays are antibody microarrays manufactured by the Felgner lab[4]. These arrays contain 500 *Plasmodium falciparum* and *P. vivax* specific antigens referred to as test probes, as well as multiple control probes (Table 2). In this study, 429 samples from 290 individuals were collected across three sites in malaria endemic regions of Zambia and Zimbabwe. Across all three study sites, a random stratified sampling scheme was employed for household selection and every individual present in the household at the time of visit was eligible for enrollment meaning that not all individuals in the study had malaria. For each individual enrolled in the study, at least one serum sample was collected and spotted on an array. Additionally, protein microarrays were spotted with sera from adults residing in the USA who had never traveled to a malaria endemic region (control serum).[7] The full content of these arrays is available in the Gene Expression Omnibus database under accession number GPL18316, and raw data has been uploaded to `GitHub (https://github.com/sberube3/pre_processing_proteomics)`.

**Table S 2:** Probe composition on the malaria array.

| Control Type | Protein | Number of Spots |
|---|---|---|
| Negative Control | Blank1 | 4 |
| | Blank2 | 5 |
| | Empty | 4 |
| | No DNA Reaction | 24 |
| | TTBS | 32 |
| Positive Control | anti-human IgG 0.003 | 4 |
| | anti-human IgG 0.03 | 4 |
| | anti-human IgG 0.3 | 4 |
| | anti-human IgG 0.001 | 4 |
| | anti-human IgG 0.01 | 4 |
| | anti-human IgG 0.1 | 4 |
| | IgG mix 0.003 | 4 |
| | IgG mix 0.03 | 4 |
| | IgG mix 0.3 | 4 |
| | IgG mix 0.001 | 4 |
| | IgG mix 0.01 | 4 |
| | IgG mix 0.1 | 4 |
| Active Proteins | Active Proteins | 1038 |

### 6 Simulation

We generate measurements that mimic the malaria arrays and the HuProt$^{TM}$ arrays in four different simulation scenarios. Each simulation scenarios compares measurements across 5 generated malaria arrays and 5 generated HuProt$^{TM}$ arrays. Generated measurements are denoted by $X_{ijkl}$, where $i$ is the array type, HuProt$^{TM}$ (H) or malaria (M), $j$ is the simulation scenario $1, 2, 3$ or $4$, $k$ is the array number A,B,C,D or E, and $l$ is the type of probe, test (T) or control (C). We further consider a subscript $C_p$ where $p$ is the type of control spot, $p \in \{1, ..., 15\}$ for the malaria arrays (M) and $p \in \{1, ..., 17\}$ for the HuProt$^{TM}$ arrays (H).

The set of generated malaria control probe measurements $X_{MjkC_p}$ contains $108$ individual measurements, drawn from $15$ different continuous uniform distributions $(U_1, ..., U_{15})$. The set of generated HuProt$TM$ control probe measurements $X_{HjkC_p}$ contains $4656$ individual measurements. These measurements are drawn from $17$ continuous uniform distributions $(U_1, ..., U_{17})$. The bounds of the uniform distributions vary depending on the scenario $j$ and the array number $k$.

In scenario $1$, the set of control probes $X_{M1kC_p}$ and $X_{H1kC_p}$ are drawn from the same uniform distributions across all five arrays A,B,C,D, and E, and the set of $1000$ malaria test probes $X_{M1kT}$ and $10000$ HuProt$^{TM}$ test probes $X_{H1kT}$ are drawn from

$$\begin{aligned} X_{i1kT} &= S \cdot e, \quad e = \exp\{Z\} \\ Z = \Phi_{0,1}^{-1}(B), \quad B &\sim \text{beta}(2, 2), \quad S \sim \text{log-normal}(\mu_S = 0, \sigma_S^2 = 1.5) \end{aligned} \tag{8}$$

across all five arrays A,B,C,D, and E. In scenario 2, the set of control probes $X_{M2kC_p}$ and $X_{H2kC_p}$ are drawn from different uniform distributions across each of the five arrays A,B,C,D, and E, (the uniform distributions from scenario 1 are multiplied by a different constant for each of the arrays B,C,D and E) and the set of 1000 malaria test probes $X_{M2kT}$ and 10000 HuProt$^{TM}$ test probes $X_{H2kT}$ are drawn from

$$\begin{aligned} X_{i2kT} &= S \cdot e, \quad e = \exp\{Z\} \\ Z = \Phi_{0,1}^{-1}(B), \quad B &\sim \text{beta}(2,2), \quad S \sim \text{log-normal}(\mu_S, \sigma_S^2) \end{aligned} \tag{9}$$

where $\mu_S = 0$ and $\sigma_S^2 = 1.5$ for Array A, $\mu_S = 0.25$ and $\sigma_S^2 = 2$ for Array B, $\mu_S = 0.5$ and $\sigma_S^2 = 2$ for Array C, $\mu_S = 0.75$ and $\sigma_S^2 = 2$ for Array D, and $\mu_S = 1$ and $\sigma_S^2 = 2$ for Array E. In scenario 3, the set of control probes $X_{M3kC_p}$ and $X_{H3kC_p}$ are drawn in the same way as scenario 1 and the set of 1000 malaria test probes $X_{M1kT}$ and 10000 HuProt$^{TM}$ test probes $X_{H1kT}$ are drawn from

$$\begin{aligned} X_{i3kT} &= S^2 \cdot e, \quad e = \exp^Z \\ Z = \Phi_{0,1}^{-1}(B), \quad B &\sim \text{beta}(2,2), \quad S \sim \text{normal}(\mu_S = 0, \sigma_S^2 = 1.5) \end{aligned} \tag{10}$$

across all five arrays A,B,C,D, and E. Finally, in Scenario 4, the set of control probes $X_{M3kC_p}$ and $X_{H3kC_p}$ are drawn in the same way as scenario 2 and the set of 1000 malaria test probes $X_{M1kT}$ and 10000 HuProt$^{TM}$ test probes $X_{H1kT}$ are drawn from

$$\begin{aligned} X_{i3kT} &= S^2 \cdot e, \quad e = \exp^Z \\ Z = \Phi_{0,1}^{-1}(B), \quad B &\sim \text{beta}(2,2), \quad S \sim \text{normal}(\mu_S, \sigma_S^2) \end{aligned} \tag{11}$$

where $\mu_S$ and $\sigma_S^2$ are the same as in scenario 2 for arrays A, B, C, D and E. In each simulation scenario, for the malaria and HuProt$^{TM}$ we coerce 12 randomly selected test probes probes to have exactly the same value for arrays A, B, C, D and E, these 12 probes serve as spike-in controls. With the generated values $X_{ijkl}$ we perform log transformation, remove fixed effects estimated by the robust linear model outlined in Section 1.2 and standardize as described in Section 1.3. We also perform a simulation with two alternative standardization methods using the median and IQR of each array and using the Winsorized sample mean and standard deviation as described in Section 1.3.

To measure the degree to which our pipeline removes technical variation, we measure mean squared error (MSE) defined as $(\bar{X}_{ijkT} - \bar{X}_{ijk'T})^2$ for $k$ not equal to $k'$, we report the sample mean and sample variance over all $k$ not equal to $k'$ (or all possible pairwise differences across arrays A, B, C, D and E) of the MSE. We also reported the mean difference of the 12 spike-in probes across all five arrays.

### 7 Between Array Technical Variation

Comparing the results of standardization with median and $\frac{\text{IQR}}{1.35}$ and standardization with Winsorized mean and standard deviation to those of traditional standardization, shows nearly identical Bland-Altman plots in the malaria arrays and for some pairs of arrays such as $(1,2)$ and $(1,3)$ from the cancer study, the traditional method of standardization appears to slightly outperform the other two methods (Figures S4, S5, S6, and Table S4).

**Table S 3:** Intercepts and slopes ($\hat{\beta}_0$ and $\hat{\beta}_1$) of OLS regression on between-array Bland-Altman plots. Sample mean difference ($\mu$) and sample concordance correlation ($\hat{\rho}_c$) of pairs of measurements across arrays.

| Arrays | Method | $\hat{\beta}_0$ | $\hat{\beta}_1$ | $\hat{\mu}$ | $\hat{\rho}_c$ |
|---|---|---|---|---|---|
| 498, 500 | RLM | 0.086 | 0.194 | 0.033 | 0.618 |
| | RLM+Std. | 0.107 | 0.215 | 0.095 | 0.606 |
| 498, 504 | RLM | -1.530 | -0.518 | -1.892 | 0.160 |
| | RLM+Std. | -0.828 | -0.525 | -1.112 | 0.196 |
| 500, 504 | RLM | -1.421 | -0.740 | -1.920 | 0.117 |
| | RLM+ Std. | -0.821 | -0.768 | -1.210 | 0.132 |
| 1,2 | RLM | 0.438 | 0.416 | 0.560 | 0.323 |
| | RLM+ Std. | 0.739 | 0.358 | 0.939 | 0.342 |
| 1,3 | RLM | 0.533 | 0.477 | 0.560 | 0.240 |
| | RLM+ Std. | 0.916 | 0.365 | 0.939 | 0.264 |
| 2,3 | RLM | 0.092 | 0.038 | 0.091 | 0.841 |
| | RLM+Std. | 0.153 | -0.007 | 0.153 | 0.847 |

Breaking down the separate goals of homogenizing both the mean and variance across arrays, a comparison of the slope, intercept and mean difference in measurements in Table ?? across malaria arrays suggests that the standardization procedure did little to homogenize the variance of measurements: the slopes are similar for RLM corrected measurements and standardized measurements. However, standardization did successfully homogenize mean values of fluorescent intensities: the intercepts and mean differences are closer to $0$ in the standardized measurements than in the RLM corrected measurements. In the cancer arrays, the slopes in Table ?? suggest that the standardization succeeded in homogenizing the variance across control serum arrays. In all cases though, the intercept and the mean differences are further from zero for the standardized values than for the RLM corrected values and in both cases, even after standardization, there is still between-array variation.

An analysis of Bland-Altman plots in Figure 3 further reveals a strong positive correlation between the mean and variance of measurements and an edge effect, as statistical features of technical variation. Both trends are more pronounced in the malaria arrays than in the cancer arrays. While additional normalization steps and analytic methods can account for the between-array technical variation remaining after standardization, some arrays, such as Arrays 1 and 504, are clear outliers and may need to be discarded from future analysis.

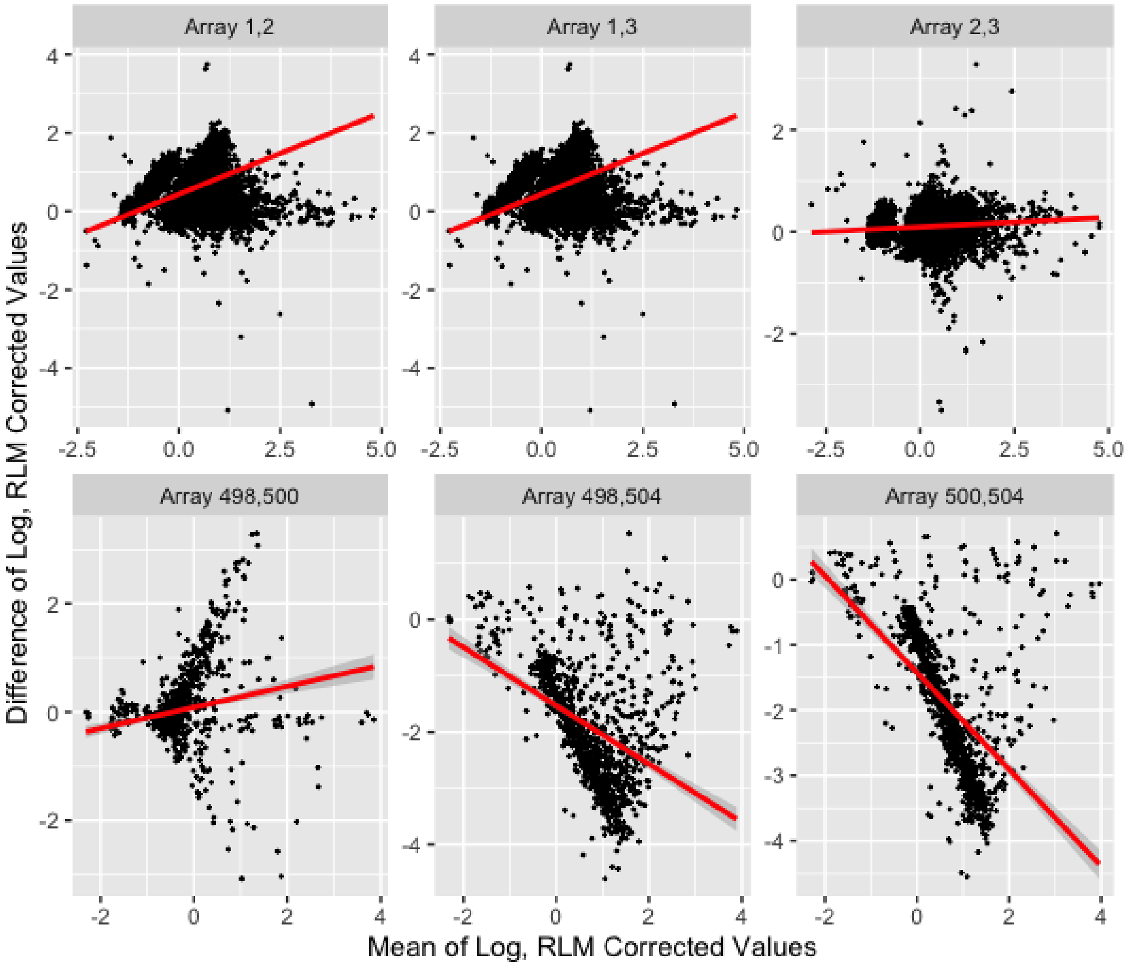


**Fig S 3:** Bland-Altman Plots of HuProt$^{TM}$ arrays spotted with cancer-free samples[8] (Arrays (1,2), (1,3), and (2,3)) and malaria arrays spotted with samples from U.S. residents[7] (Arrays (498,500),(498,504) and (500,504)). We show $M$ vs $D$ with $\tilde{Y}$s from pairs of arrays. The ordinary least squares line is shown in red.

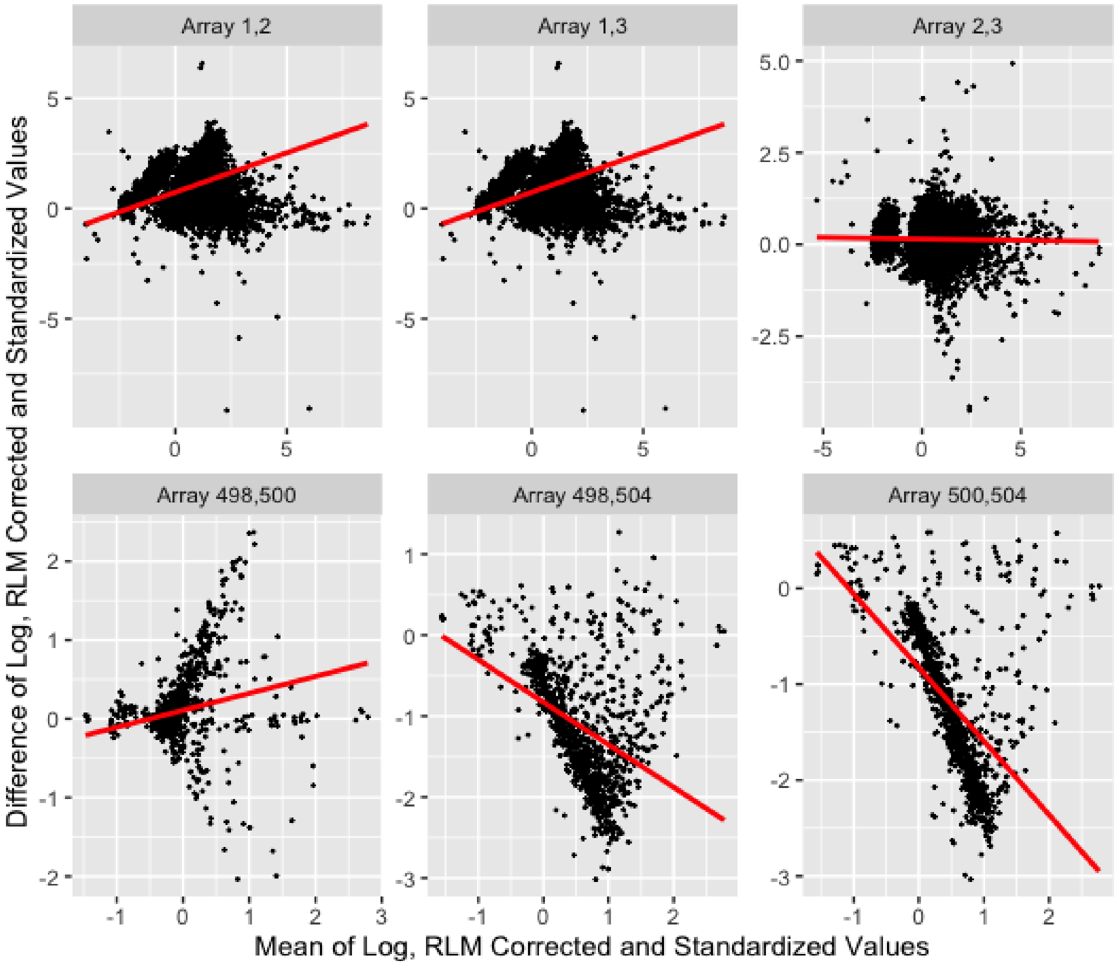


**Fig S 4:** Bland-Altman Plots of arrays spotted with cancer-free samples (Arrays (1,2), (1,3), and (2,3)) [8] and U.S. residents (Arrays (498,500),(498,504) and (500,504)) [7]. We show $M$ vs $D$ with $Y$s from pairs of arrays. The ordinary least squares line is shown in red.

### 8 Simulation Results

Generally, traditional standardization with mean and standard deviation outperforms standardization with median and $\frac{\text{IQR}}{1.35}$, and standardization with Winsorized means and standard deviations. While there are certain scenarios (Scenarios 2 and 4) where standardization with the median and IQR outperforms traditional standardization for the malaria arrays, the traditional method of standardization outperforms the two other methods in all four scenarios for the HuProt$^{TM}$ arrays (Table 5).

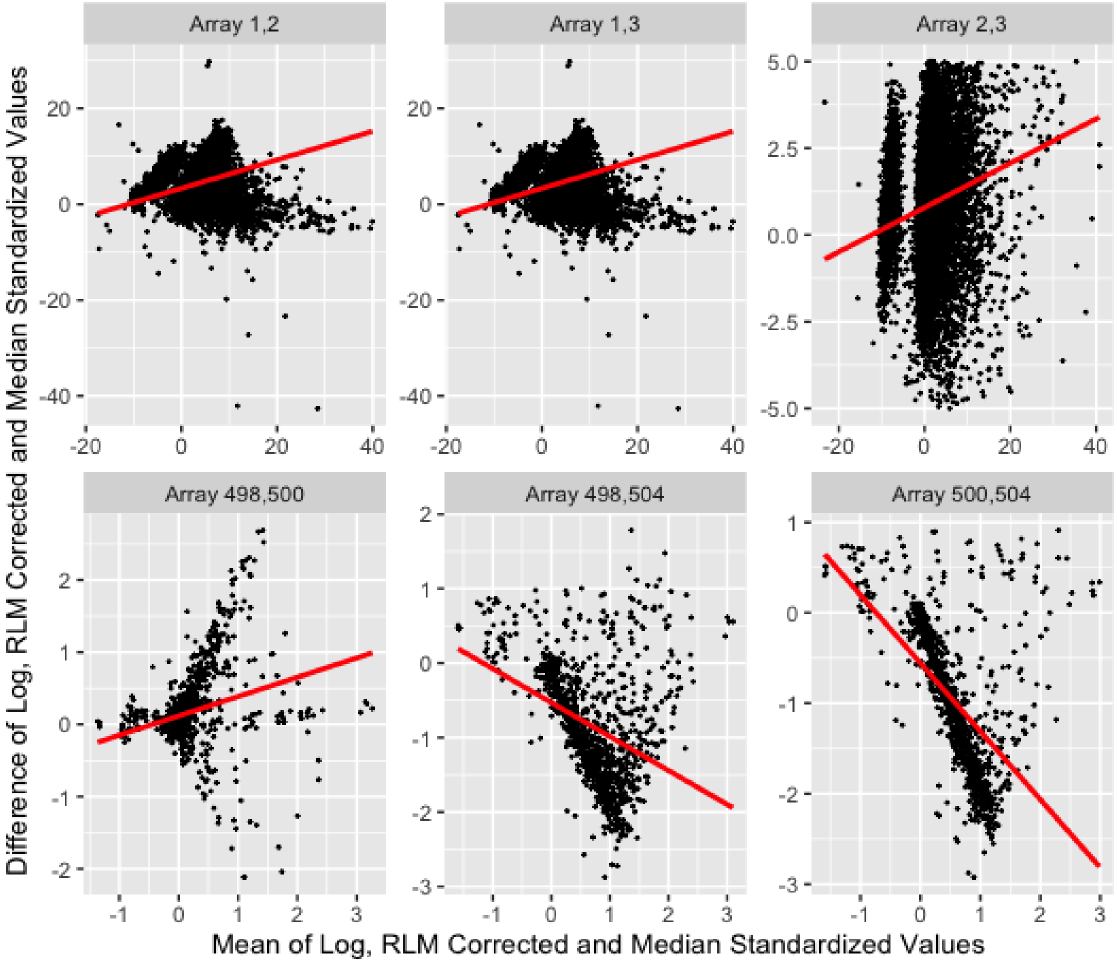


**Fig S 5:** Bland-Altman Plots of arrays spotted with cancer-free samples(Arrays (1,2), (1,3), and (2,3)) [8] and U.S. residents (Arrays (498,500),(498,504) and (500,504)) [7]. We show $M$ vs $D$ with $Y$s standardized using the median and $\frac{\text{IQR}}{1.35}$ from pairs of arrays. The ordinary least squares line is shown in red.

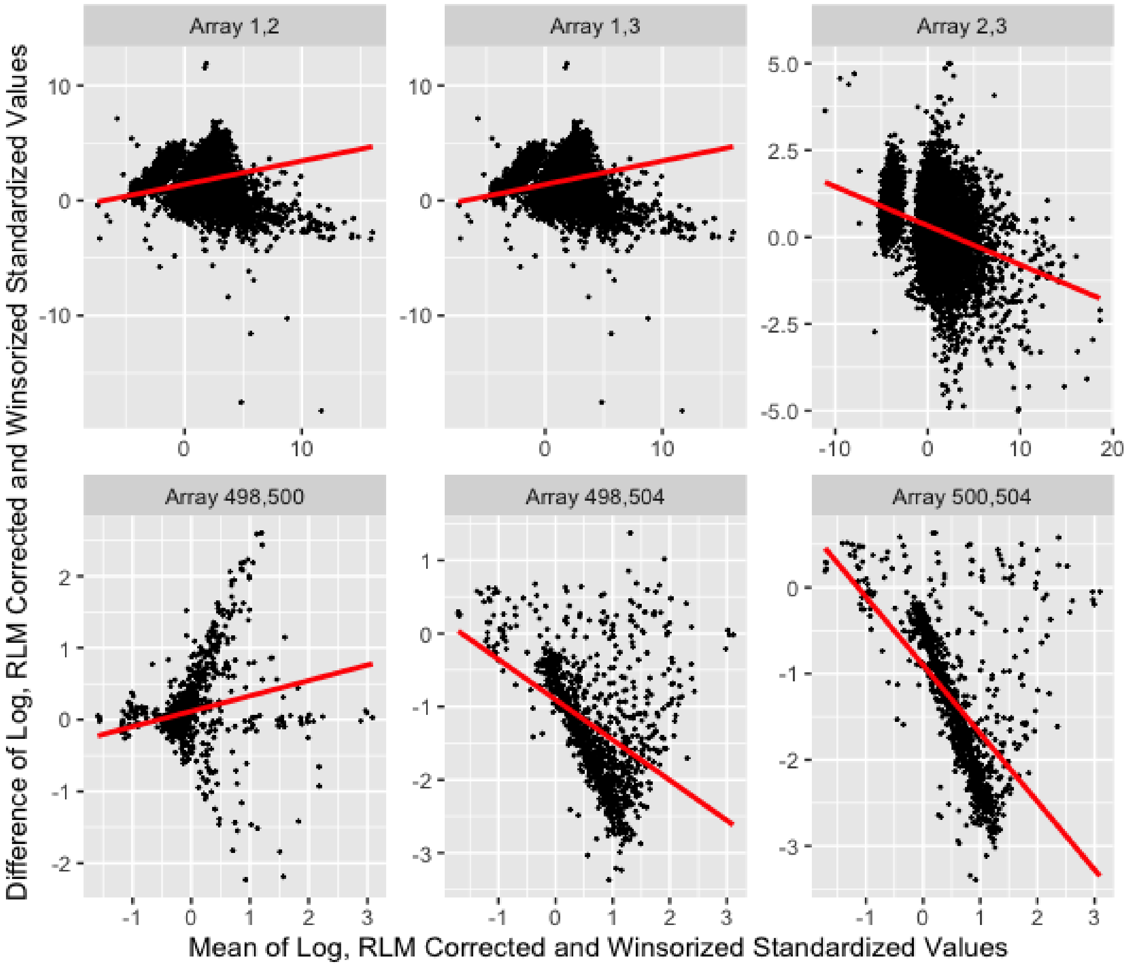


**Fig S 6:** Bland-Altman Plots of arrays spotted with cancer-free samples(Arrays (1,2), (1,3), and (2,3)) [8] and U.S. residents(Arrays (498,500),(498,504) and (500,504)) Kobayashi et al. [7] . We show $M$ vs $D$ with $Y$s standardized using the Winsorized mean and standard deviation (bottom and top 5% of data points replaced) from pairs of arrays. The ordinary least squares line is shown in red.

**Table S 4:** Intercepts and slopes ($\hat{\beta}_0$ and $\hat{\beta}_1$) of OLS regression on between-array Bland-Altman plots. Sample mean difference ($\mu$) and sample concordance correlation ($\hat{\rho}_c$) of pairs of measurements across arrays.

| Arrays | Method | $\hat{\beta}_0$ | $\hat{\beta}_1$ | $\hat{\mu}$ | $\hat{\rho}_c$ |
|---|---|---|---|---|---|
| 498, 500 | Std Mean+SD | 0.107 | 0.215 | 0.095 | 0.606 |
| | Std Med +IQR | 0.118 | 0.267 | 0.162 | 0.586 |
| | Std Wins Mean+SD | 0.116 | 0.214 | 0.107 | 0.605 |
| 498, 504 | Std Mean+SD | -0.828 | -0.525 | -1.11 | 0.196 |
| | Std Med+IQR | -0.531 | -0.456 | -0.833 | 0.281 |
| | Std Wins Mean+SD | -0.905 | -0.551 | -1.26 | 0.192 |
| 500, 504 | Std Mean+SD | -0.821 | -0.768 | -1.21 | 0.132 |
| | Std Med+IQR | -0.557 | -0.754 | -0.996 | 0.176 |
| | Std Wins Mean+SD | -0.901 | -0.792 | -1.36 | 0.129 |
| 1,2 | Std Mean+SD | 0.739 | 0.358 | 0.939 | 0.342 |
| | Std Med+IQR | 3.32 | 0.297 | 4.26 | 0.345 |
| | Std Wins Mean+SD | 1.39 | 0.205 | 1.59 | 0.374 |
| 1,3 | Std Mean+SD | 0.916 | 0.365 | 0.939 | 0.264 |
| | Std Med+IQR | 4.03 | 0.390 | 4.26 | 0.246 |
| | Std Wins Mean+SD | 1.85 | 0.075 | 1.59 | 0.299 |
| 2,3 | Std Mean+SD | 0.153 | -0.007 | 0.153 | 0.847 |
| | Std Med+IQR | 0.797 | 0.070 | 0.893 | 0.834 |
| | Std Wins Mean+SD | 0.325 | -0.117 | 0.322 | 0.842 |

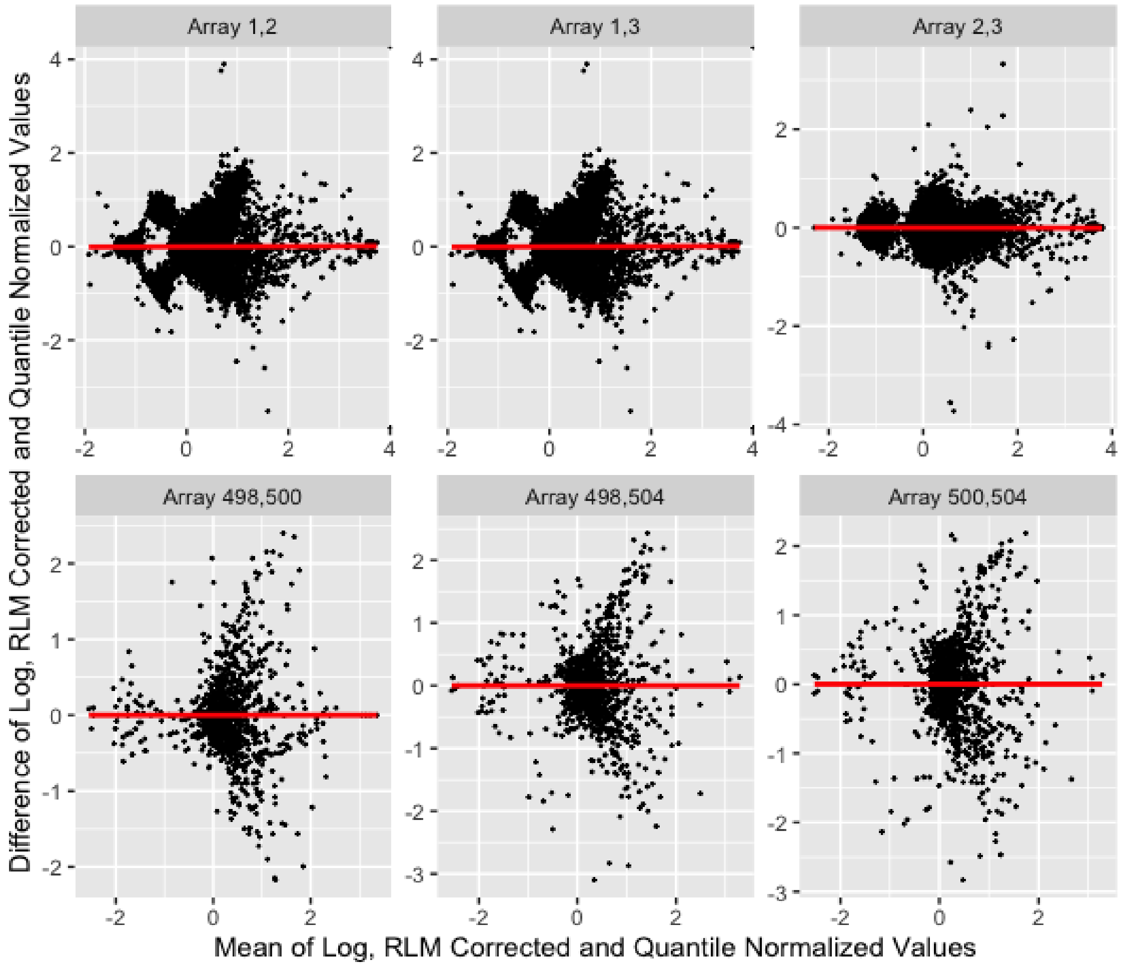


**Fig S 7:** Bland-Altman Plots of arrays spotted with cancer-free samples (Arrays (1,2), (1,3), and (2,3))[8] and U.S. residents(Arrays (498,500),(498,504) and (500,504))[7] . We show $M$ vs $D$ with values normalized using the quantile method[3] from pairs of arrays. The ordinary least squares line is shown in red.

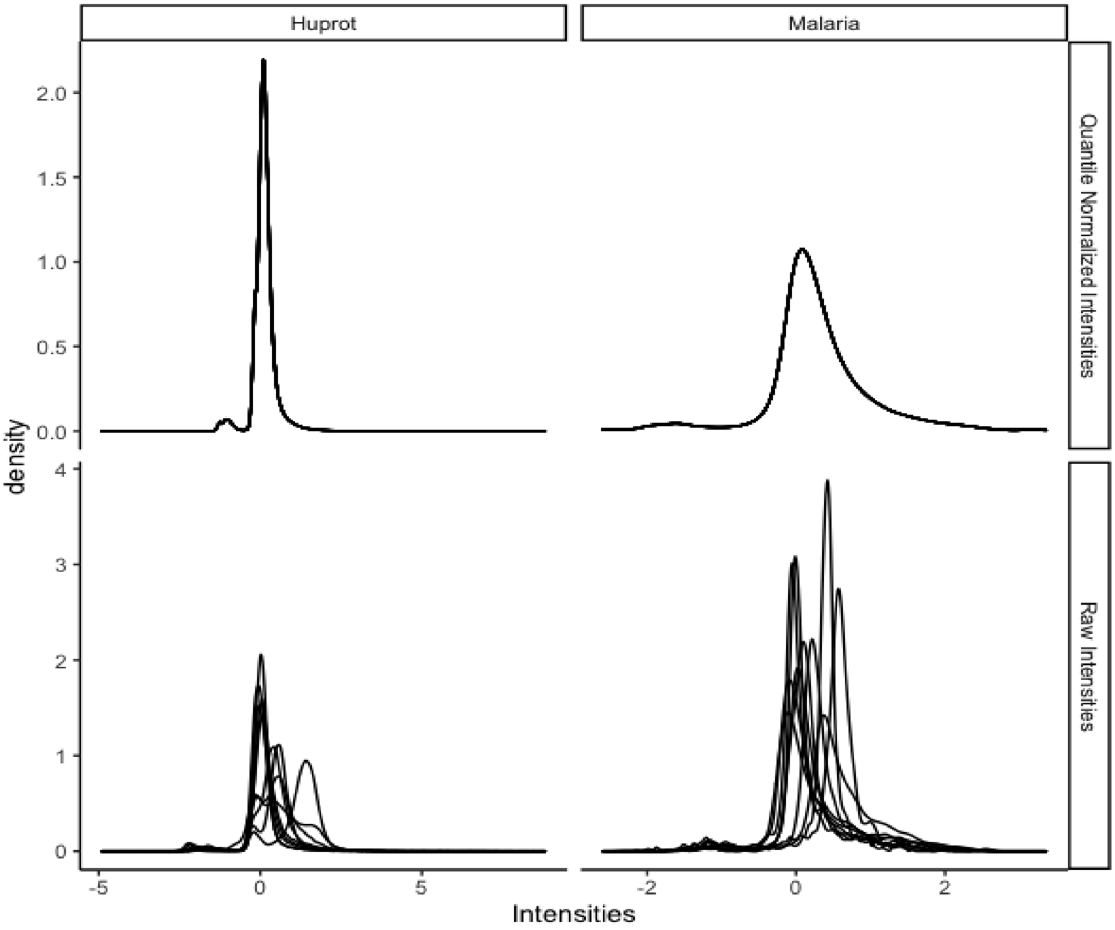


**Fig S 8:** Density of intensities from 10 randomly selected arrays from each study for values standardized values, $Y$, and for for quantile normalized values.

**Table S 5:** Mean squared error (MSE) of difference in means and variances across pairs of arrays, and mean differences of spike-in controls across arrays. Sample mean of 1000 simulations and, sample standard deviation. Scenario 1: data generating mechanism with a log-normal distribution and no difference in distributions across arrays. Scenario 2: data generating mechanism with a log-normal distribution and a difference in distributions across arrays. Scenario 3: data generating mechanism with a normal distribution and squared values and no difference in distributions across arrays. Scenario 4: data generating mechanism with a normal distribution and squared values and a difference in distributions across arrays.

| Array Type | Scenario | Method | MSE Mean | MSE Var | Mean Difference |
|---|---|---|---|---|---|
| Malaria | 1 | Std Mean+SD | $0.027_{0.018}$ | $0.055_{0.040}$ | $-0.001_{0.033}$ |
| | | Std Med+IQR | $1.365_{0.225}$ | $7.186_{1.045}$ | $0.001_{0.046}$ |
| | | Std Wins Mean+SD | $0.037_{0.024}$ | $0.092_{0.067}$ | $0.001_{0.044}$ |
| | 2 | Std Mean+SD | $0.151_{0.062}$ | $2.950_{0.494}$ | $0.356_{0.035}$ |
| | | Std Med+IQR | $0.127_{0.060}$ | $1.823_{0.433}$ | $0.312_{0.051}$ |
| | | Std Wins Mean+SD | $0.184_{0.080}$ | $4.148_{0.757}$ | $0.387_{0.046}$ |
| | 3 | Std Mean+SD | $0.020_{0.015}$ | $0.053_{0.039}$ | $0.001_{0.038}$ |
| | | Std Med+IQR | $0.031_{0.022}$ | $0.052_{0.037}$ | $0.003_{0.055}$ |
| | | Std Wins Mean+SD | $0.031_{0.023}$ | $0.081_{0.060}$ | $0.001_{0.051}$ |
| | 4 | Std Mean+SD | $0.172_{0.059}$ | $0.219_{0.115}$ | $0.291_{0.037}$ |
| | | Std Med+IQR | $0.103_{0.054}$ | $0.174_{0.099}$ | $0.237_{0.060}$ |
| | | Std Wins Mean+SD | $0.200_{0.075}$ | $0.320_{0.172}$ | $0.312_{0.050}$ |
| HuProt$^{TM}$ | 1 | Std Mean+SD | $0.006_{0.004}$ | $0.215_{0.153}$ | $-0.001_{0.014}$ |
| | | Std Med+IQR | $0.015_{0.011}$ | $1.368_{0.988}$ | $-0.001_{0.027}$ |
| | | Std Wins Mean+SD | $0.017_{0.011}$ | $1.235_{0.958}$ | $-0.001_{0.019}$ |
| | 2 | Std Mean+SD | $0.293_{0.043}$ | $33.703_{3.337}$ | $0.771_{0.016}$ |
| | | Std Med+IQR | $0.593_{0.098}$ | $138.530_{16.448}$ | $1.096_{0.032}$ |
| | | Std Wins Mean+SD | $0.827_{0.120}$ | $266.116_{21.497}$ | $1.295_{0.026}$ |
| | 3 | STD Mean+SD | $0.004_{0.003}$ | $0.110_{0.078}$ | $-0.000_{0.018}$ |
| | | Std Med+IQR | $0.013_{0.009}$ | $0.669_{0.490}$ | $-0.001_{0.035}$ |
| | | Std Wins Mean+SD | $0.011_{0.008}$ | $0.760_{0.564}$ | $-0.000_{0.028}$ |
| | 4 | Std Mean+SD | $1.758_{0.118}$ | $17.511_{2.488}$ | $0.826_{0.040}$ |
| | | Std Med+IQR | $5.058_{0.345}$ | $257.858_{28.043}$ | $1.215_{0.077}$ |
| | | Std Wins Mean+SD | $8.492_{0.427}$ | $603.182_{46.494}$ | $1.499_{0.086}$ |

**Table S 6:** Mean squared error (MSE) of difference in means and variances across pairs of arrays, and mean differences of spike-in controls across arrays. Sample mean of 1000 simulations and, in subscript, the sample standard deviation.

| Array Type | Scenario | Method | MSE Mean | MSE Var | Mean Difference |
|---|---|---|---|---|---|
| Malaria | 1 | RLM+ Std. | $0.003_{0.002}$ | $0.006_{0.004}$ | $0.001_{0.033}$ |
| | | Quantile | $0.000_{0.000}$ | $0.000_{0.000}$ | $0.001_{0.004}$ |
| | | VSN | $0.000_{0.000}$ | $0.001_{0.000}$ | $0.000_{0.000}$ |
| | | CL | $0.004_{0.000}$ | $0.108_{0.006}$ | $-0.000_{0.005}$ |
| | 2 | RLM+Std. | $0.015_{0.006}$ | $0.296_{0.047}$ | $0.357_{0.035}$ |
| | | Quantile | $0.000_{0.000}$ | $0.000_{0.000}$ | $0.215_{0.007}$ |
| | | VSN | $0.000_{0.000}$ | $0.000_{0.000}$ | $0.026_{0.000}$ |
| | | CL | $0.007_{0.000}$ | $4.631_{0.855}$ | $0.221_{0.014}$ |
| | 3 | RLM+ Std. | $0.002_{0.001}$ | $0.005_{0.004}$ | $-0.001_{0.037}$ |
| | | Quantile | $0.000_{0.000}$ | $0.000_{0.000}$ | $0.005_{0.001}$ |
| | | VSN | $0.002_{0.000}$ | $0.066_{0.071}$ | $-0.001_{0.001}$ |
| | | CL | $0.027_{0.000}$ | $3.766_{8.296}$ | $0.005_{0.103}$ |
| | 4 | RLM+Std. | $0.018_{0.006}$ | $0.022_{0.012}$ | $0.291_{0.037}$ |
| | | Quantile | $0.000_{0.000}$ | $0.000_{0.000}$ | $0.589_{0.014}$ |
| | | VSN | $0.007_{0.000}$ | $0.255_{1.421}$ | $0.199_{0.001}$ |
| | | CL | $0.027_{0.000}$ | $3.844_{8.540}$ | $0.668_{0.104}$ |
| HuProt | 1 | RLM+ Std. | $0.003_{0.002}$ | $0.006_{0.004}$ | $0.001_{0.033}$ |
| | | Quantile | $0.000_{0.000}$ | $0.000_{0.000}$ | $0.001_{0.000}$ |
| | | VSN | $0.000_{0.000}$ | $0.001_{0.000}$ | $0.000_{0.000}$ |
| | | CL | $0.001_{0.000}$ | $0.007_{0.000}$ | $0.000_{0.000}$ |
| | 2 | RLM+Std. | $0.015_{0.006}$ | $0.296_{0.047}$ | $0.357_{0.035}$ |
| | | Quantile | $0.000_{0.000}$ | $0.000_{0.000}$ | $0.294_{0.003}$ |
| | | VSN | $0.007_{0.000}$ | $0.012_{0.000}$ | $0.320_{0.001}$ |
| | | CL | $0.004_{0.000}$ | $2.538_{0.032}$ | $0.315_{0.009}$ |
| | 3 | RLM+ Std. | $0.002_{0.001}$ | $0.005_{0.004}$ | $-0.001_{0.037}$ |
| | | Quantile | $0.000_{0.000}$ | $0.000_{0.000}$ | $0.001_{0.001}$ |
| | | VSN | $0.000_{0.000}$ | $0.001_{0.000}$ | $-0.000_{0.000}$ |
| | | CL | $0.001_{0.000}$ | $0.248_{0.032}$ | $-0.008_{0.113}$ |
| | 4 | RLM+Std. | $0.018_{0.006}$ | $0.022_{0.012}$ | $0.291_{0.037}$ |
| | | Quantile | $0.000_{0.000}$ | $0.000_{0.000}$ | $0.670_{0.001}$ |
| | | VSN | $0.011_{0.000}$ | $0.068_{0.000}$ | $0.467_{0.000}$ |
| | | CL | $0.019_{0.000}$ | $0.348_{0.055}$ | $0.799_{0.134}$ |

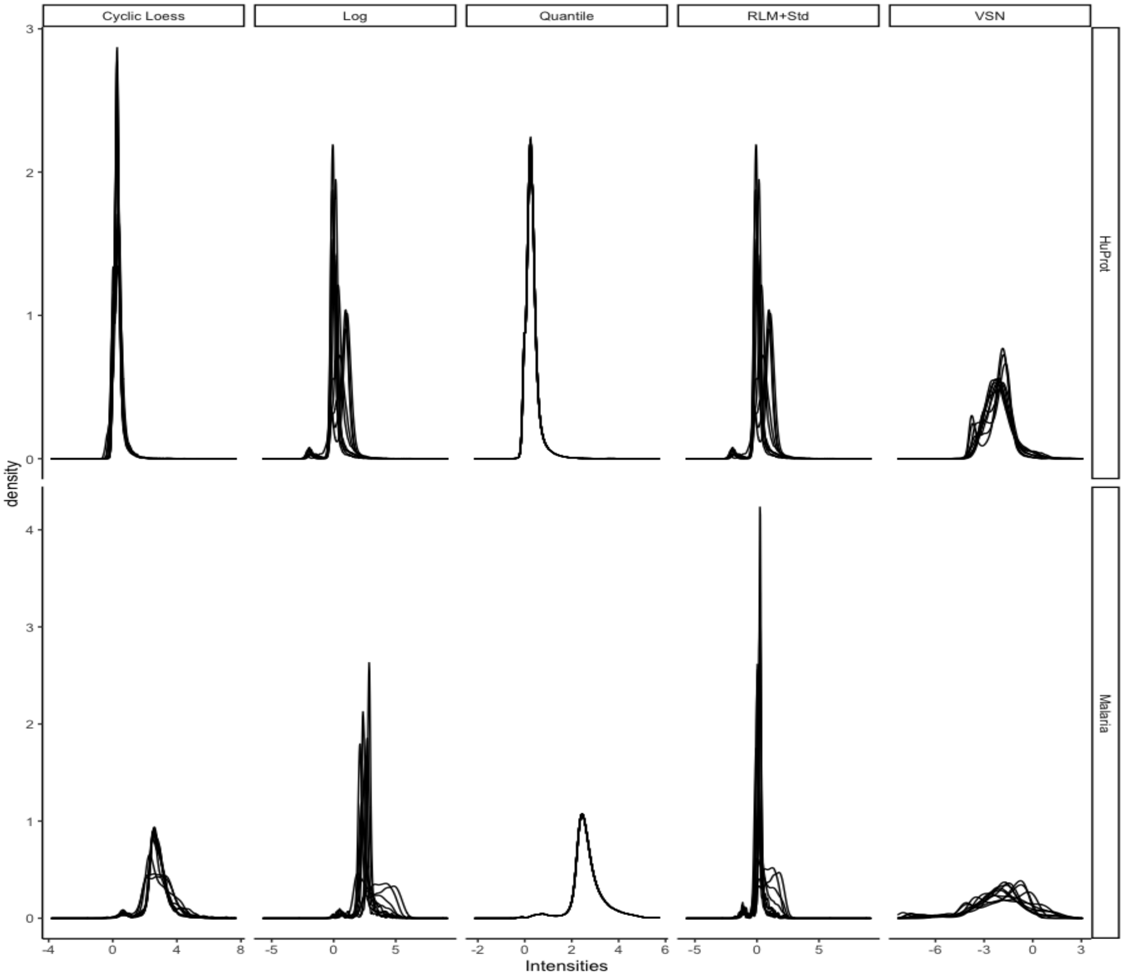


**Fig S 9:** Densities of fluorescent intensities on $10$ randomly chosen arrays from the malaria study and the lung cancer study after different normalization methods. Chosen arrays are spotted with sera from malaria endemic countries in the malaria study and individuals with lung cancer in the lung cancer study (non-control sera arrays).
